# Lactoferrin Reverses Methotrexate Driven Epithelial Barrier Defect by Inhibiting TGF-β Mediated Epithelial to Mesenchymal Transition

**DOI:** 10.1101/2019.12.23.878207

**Authors:** Thomas E. Wallach, Vasudha Srivastava, Efren Reyes, Ophir D. Klein, Zev J. Gartner

**Author notes:** CORRESPONDENCE: Zev J. Gartner, 600 16th St. Rm N512E, UCSF Box 2280, San Francisco, CA 94158, United States. DISCLOSURES: None of the authors have financial disclosures. ABBREVIATIONS: Methotrexate (MTX), Lactoferrin (LTF), Epithelial to Mesenchymal Transition (EMT). AUTHOR CONTRIBUTIONS: T.E.W. and Z.J.G. conceptualized the study. T.E.W. designed experiments, carried out barrier function experiments in Caco2 and enteroid models, RNA sequencing analysis, drafted the manuscript, and figure design. V.S. designed experiments, performed RNAseq analysis, figure design, statistical analyses, and edited the manuscript. O.D.K. provided material support for enteroid culture and edited the manuscript. E.R. assisted in enteroid culture, experimental design, imaging, and co-wrote the manuscript. Z.J.G. designed experiments, supervised the research, and co-wrote and edited the manuscript.

## Abstract

**BACKGROUND AND AIMS:** Methotrexate is an important tool in the arsenal of oncologists, gastroenterologists, and rheumatologists. At low doses it induces intestinal barrier dysfunction that may induce side effects such as gastrointestinal discomfort and liver injury. Previous studies suggest that lactoferrin can improve barrier function in a variety of contexts. This study set out to determine the mechanism of methotrexate induced barrier dysfunction and assess the effect of lactoferrin and other components of human breast milk on this dysfunction.

**METHODS:** Using a murine enteroid model and Caco2 spheroids, we measured flux of basolateral-administered fluorescent dextran into the lumen. Barrier dysfunction was induced using methotrexate (220 nM) or lipopolysaccharide (20 nM). Human lactoferrin was added at 0.8 mg/ml (10 µM). RNAseq was performed on exposed samples.

**RESULTS:** Lactoferrin blocks methotrexate-induced barrier dysfunction in murine enteroids. Similar results were observed when barrier dysfunction was induced in Caco2 spheroids with methotrexate and LPS, but not ML7. RNAseq revealed activation of TGF-β response genes and epithelial-mesenchymal transition (EMT) by methotrexate, which normalized in the presence of lactoferrin. TGF-β receptor inhibition (RepSox) blocked methotrexate induced barrier dysfunction in Caco2 spheroids. 20 nM TGF-β induced barrier dysfunction in Caco2 spheroids which was also inhibited by lactoferrin.

**CONCLUSIONS:** Methotrexate induces barrier dysfunction by activation of an EMT program promoted by TGF-β signaling and inhibited by lactoferrin. Lactoferrin is also protective of barrier function in an LPS-induced model. The likely mechanism of this effect is blockade of EMT programs induced by TGF-β.

## INTRODUCTION

Intestinal barrier dysfunction is implicated in the pathology of multiple complex diseases of the gut, as well as those involving tissues outside of the gastrointestinal system. Celiac disease^1^, fatty liver disease^2^, Irritable Bowel Syndrome (IBS) and functional abdominal disorders^3^, Inflammatory Bowel Syndrome (IBD), and metabolic syndrome^4^ have all been tied to alterations in intestinal epithelial function in the developed world. In the developing world, barrier function has been implicated in patients with environmental enteropathy^5^, a condition causing stunting and increased risk of metabolic syndrome that affects hundreds of millions of children in the developing world^5^. In addition to its contribution to gastrointestinal disease, intestinal barrier dysfunction may contribute to medication-limiting side effects^6^. Methotrexate is an illustrative example of such a pharmacological agent. As part of the World Health Organization’s list of Essential Medicines, it is a critically important drug that is commonly used to treat many forms of cancer and autoimmune disease. However, methotrexate is known to induce significant side effects, with abdominal discomfort and potential liver injury even at low doses, and mucositis at doses used in oncology^7^. Given the known barrier dysfunction caused by methotrexate^8–10^, it is possible that barrier protective substances could mitigate these dose limiting effects.

Development and broad administration of new therapeutics capable of modulating intestinal barrier function could dramatically expand the utility of Methotrexate, and could also have a major impact on the variety of other diseases linked to barrier dysfunction. While several compounds have been shown to decrease barrier function fewer exist that increase barrier function. Several potential strategies exist for developing such drugs to counteract the effects of methotrexate. For example, compounds which target the specific mechanism through which methotrexate induces barrier dysfunction could be developed, so long as the mechanism or target tissue is distinct from that which methotrexate elicits its therapeutic activity. Alternatively, compounds could be developed that act locally in the gut to inhibit the effects of methotrexate on the intestinal epithelium, but which are not absorbed significantly and therefore cannot act on more distal sites. Independent of their mechanism of action, such compounds must additionally be of unusually low cost and high safety in order to justify their use. Thus, agents capable of modifying barrier function that are readily available, inexpensive, and generally regarded as safe (GRAS)^11^ would provide ideal therapeutics to counteract the effects of methotrexate on the intestinal epithelium.

The component of human breast milk (HBM) satisfy many of these criteria. They are core components of the early human diet and have been shown conclusively to improve gastrointestinal and metabolic outcomes^12, 13^. The precise mechanism by which breast milk impacts gastrointestinal function is not known, but it has been hypothesized to act primarily through modulation of the microbiota – for example, by providing unique metabolic substrates or through the antimicrobial properties of some HBM components. Among the constituents of milk, lactoferrin has received particular attention because of its antimicrobial properties^14^, immune regulatory properties^15^, and ability to signal through the lactoferrin receptor expressed on epithelial cells^16^. It is an approximately 80 kDa iron binding protein that is present in breast milk at concentrations ranging from 10 to 90 mg/ml (13-113 μM)^17^, but may be deactivated during pasteurization^18^. It is also present in cow’s milk at the significantly lower concentration of 0.031-0.485 mg/ml^19^. Taken orally, it has been shown that approximately 60-70% of lactoferrin survives gastric hydrolysis^20^ to enter the intestine, with likely higher ability to tolerate the stomach in neonates due to lower levels of gastric acidity^21^. Numerous clinical studies indirectly suggest a more generalized capacity of lactoferrin to reinforce the body’s epithelial barriers, such as a capacity to decrease incidence of necrotizing enterocolitis (NEC) in premature infants^22^, decrease days and severity of diarrheal illness^23–25^, and protect against upper respiratory tract infection^23^. In addition, lactoferrin has demonstrated the capacity to protect from methotrexate induced barrier dysfunction in rats^26^. Particularly intriguing is a recent report that suggests that lactoferrin improves barrier function in the context of environmental enteropathy^27^. Despite the multiple activities of lactoferrin in the gastrointestinal tract, how lactoferrin specifically acts to improve barrier function is unclear. Lactoferrin’s effects on the epithelium could be indirect – that is downstream of changes in the immune system or microbiota – or direct, immediately downstream of its activity on the epithelial lactoferrin receptor or other molecules in the immediate epithelial microenvironment.

To determine whether lactoferrin acts directly on the epithelium to modulate epithelial barrier function, an *in vitro* model lacking the immune and microbial compartments would be useful. Previous *in vitro* studies of barrier function have primarily focused on two-dimensional culture models, assessing impact via trans-epithelial electrical resistance (TEER) or the flux of dyes across the monolayer^28^. However, recent work has illustrated the importance of three dimensional tissue structure to barrier function and cellular differentiation, with 2D monolayers displaying drastically different behavior than 3D culture^29^. In addition, advances in tissue engineering have allowed for the culture of three-dimensional organoids derived from primary tissue samples or stem cells, allowing for more accurate modeling of organ function, in particular that of the intestinal epithelium^30–32^. In this study, we use enteroids (organoids derived from the intestinal epithelium) and 3D culture models to test the hypothesis that lactoferrin directly impacts intestinal epithelial barrier function and would be protective from barrier injury induced by methotrexate or bacterial LPS.

## MATERIALS AND METHODS

### Chemicals and reagents

Recombinant human holo-lactoferrin (iron-saturated) (L1294) was purchased from Sigma-Aldrich. Aliquots were prepared by resuspending in PBS at a concentration of 1 mg/ml and were stored at −20 °C. Methotrexate (M9929) was purchased from Sigma-Aldrich and prepared by dissolving in PBS to a concentration of 1 mg/ml. Solutions were stored at 4 °C and used within 2 days. Recombinant human TGF-β (7754-BH) was purchased from R&D Systems (Minneapolis, MI). It was reconstituted at 100 μg/mL in sterile 4 mM HCl containing at least 0.1% BSA, aliquoted, and stored at −80°C as per manufacturer product specifications. RepSox was purchased from Sigma Aldrich, and 10X stocks were prepared by suspending in DMSO at 25 mg/ml and stored at −20 °C. LPS-EK (tlrl-eklps) was obtained from Invivogen (San Diego, CA), suspended in PBS at a concentration of 1 mg/ml, and aliquots were stored at −20 °C. ML-7 was purchased from Tocris Biosciences (Bristol, United Kingdom) and resuspended in DMSO to a concentration of 50 mM and stored at −20 °C prior to use. Expressed breast milk (EBM) was obtained by anonymous donation from a G1P1 human female on month 6 of breast-feeding post-partum. Use of EBM in this study was determined to be exempt by the UCSF Institutional Review Board.

### Cell Culture

Caco2 cells (ATCC HTB-37) were obtained from UCSF cell culture facility and maintained in culture as 2D monolayers. They were maintained in DMEM medium supplemented with 20% FBS, 100 U/ml Penicillin/Streptomycin, and 1% NEAA, all purchased from Gibco (Boston, MA). Caco2 cells were passaged at 70-80% confluency using 0.5% trypsin. Cells were transduced with pSicoR-Ef1a-EGFP lentivirus to express cytosolic GFP to facilitate lumen visualization. GFP positive cells were sorted by fluorescence activated cell sorting (FACS) using BD AriaIII cell sorter (BD Biosciences), and expanded. Caco2 spheroids were cultured by resuspension of dissociated cells in Matrigel (Corning, Corning, NY) at a concentration of 5 x 10^5^ cell/ml and cultured for 6-8 days before experimental use. Passage number ranged from 28-35 for all Caco2 experiments.

Murine enteroids were established from *Vil1^Cre-ERT^*^2^*^/+^ R26^mTmG/mTmG^* mice^33, 34^ and were kindly provided by Dr. Kara Mckinley at UCSF^35^. This strain expressed membrane bound td-tomato fluorescent protein in the intestinal epithelium. Enteroids were cultured according to established protocols^32, 36^. Briefly, enteroids were passaged every 7 days by mechanical disruption and enteroid fragments were embedded in 50 µl Matrigel dome droplets on pre-warmed 24-well tissue culture plastic plates. Dome droplets were then overlaid with 500 µl of ENR medium, and medium was changed every 4 days^4, 32, 361^, as per Mahe et al^32^. For all enteroid cultures, recombinant human R-Spondin was replaced with 5% R-Spondin conditioned medium.

### In-vitro Permeability Assay

Three dimensional microtissues (Caco2 spheroids or murine enteroids) were cultured in Matrigel for 7 and 3 days respectively. The spheroid/enteroid cultures were incubated for 4 hours with fluorescently-labeled 3 kDa dextran-dye conjugates to equilibrate in Matrigel. For Caco2 experiments, a Texas-red dextran-conjugate was used (Thermo Fisher) and for enteroids a cascade blue-dextran and forest green-dextran conjugate was used (Thermo Fisher). Barrier disruption was initiated by the addition of methotrexate at a concentration of 220 nM. Lactoferrin dosage was assessed via exposure of multiple concentrations via end point analysis of luminal barrier intensity at 14 hrs. Experimental assays were performed with addition of lactoferrin 10 µM or repsox (348 nM). Live imaging was performed hourly for 12 hours using a Zeiss LSM800 with incubation at 37 °C with 5% CO_2_. TGF-β and LPS assays were performed identically, but with TGF-β in place of methotrexate at a concentration of 220 nM and LPS at a concentration of 20 nM.

### Image Analysis

Normalized luminal intensity was measured as the ratio of luminal to extraluminal average fluorescence intensity, and was quantified using Fiji/ImageJ^37^. Relative lumen intensity was defined as the change in the normalized lumen intensity compared to the start of the experiment (0 h). Dose response was performed by aggregating average endpoint readings per well, with 2-3 well replicates. 20 spheroids/enteroids were measured per experimental well. Aggregated data was analyzed using R-Suite (Warsaw, Poland). Statistical significance of difference across conditions was calculated using a non-parametric Wilcox test compared to methotrexate treatment.

### RNA Sequencing

Caco-2 spheroids were exposed to methotrexate, lactoferrin, and lactoferrin with methotrexate. RNA was extracted using RNeasy Plus Micro kit (Qiagen, Hilden, Germany) after 4 hour exposure as per protocol. The cDNA library preparation and paired-end sequencing was performed by Novogene Corporation Inc (Sacramento, CA). Analysis was performed using the DEseq2^38^ package in R-Suite (Warsaw, Poland). Differentially expressed genes were identified as genes that had >2-fold change in expression with adjusted p-value less than 0.01. Gene ontology analysis on the top 500 upregulated genes was performed using the PGSEA package in R^39^. The log-normalized read counts are provided in **Supplemental Table 1**. A list of differentially expressed genes is provided in **Supplemental Table 2**.

### Immunofluorescence

3D cultures were fixed in 4% paraformaldehyde for 45 minutes and washed 3x with PBS. Fixed tissues were permeabilized with Triton X-100 for 15 min at room temperature then blocked overnight in PBS supplemented with 1% BSA, 0.3% Triton X-100, 10% goat serum, and 1:50 goat-anti-mouse IgG Fab fragment (Jackson Immunoresearch, 115-007-003). E-cadherin was visualized with rat anti-E-cadherin (Invitrogen #13-1900), both at 1:100 concentrations as per manufacturer protocol. Actin was stained with phalloidin 405 (Invitrogen) at 1:1000 concentration. Secondary staining was performed using Invitrogen Alexa-fluor secondary antibodies at 1:100 concentration (#A-11001, A-21247). Imaging was performed using a Zeiss Observer.Z1 with a Yokogawa CSUX1 spinning disk.

### Ethics

Approval waiver was sought and obtained for use of human breast milk without need for full IRB approval.

## RESULTS

### Lactoferrin Inhibits Methotrexate Induced Barrier Dysfunction in Three dimensional Intestinal Epithelial Cell Culture Models

To establish a model of epithelial barrier dysfunction, we initially used immortalized colonic adenocarcinoma (Caco2) cells^40^. These cells form three-dimensional spheroids after plating in Matrigel and are mature with an intact barrier surrounding a hollow lumen after approximately 6 days in culture. These spheroids are amenable to quantitative assessment of barrier integrity by measuring the flux of fluorescently-labeled dextran (3 kDa) across the barrier. Methotrexate induces non-apoptotic barrier dysfunction both *in vivo* and *in vitro,*^8, 9^ and we also found that 220 nM methotrexate administered to the basolateral surface of Caco2 cysts left them structurally intact, but unable to exclude a 3 kDa dextran (Figure 1A). In this assay, we found that the luminal fluorescence of cysts gradually increased upon methotrexate administration but without significant apoptosis, indicative of a disrupted barrier (Figure 1B, Supplemental Figure S1A, B). Consistent with previous in vivo studies, concurrent addition of breast milk provided a protective effect against methotrexate treatment supporting the in vivo relevance of the permeability assay (Figure 1C). However, in this case we observe a restoration of barrier function independent of a functional microbiota or immune system. Thus, components of breast milk can act directly on the epithelium to improve barrier function.

**Figure 1:**
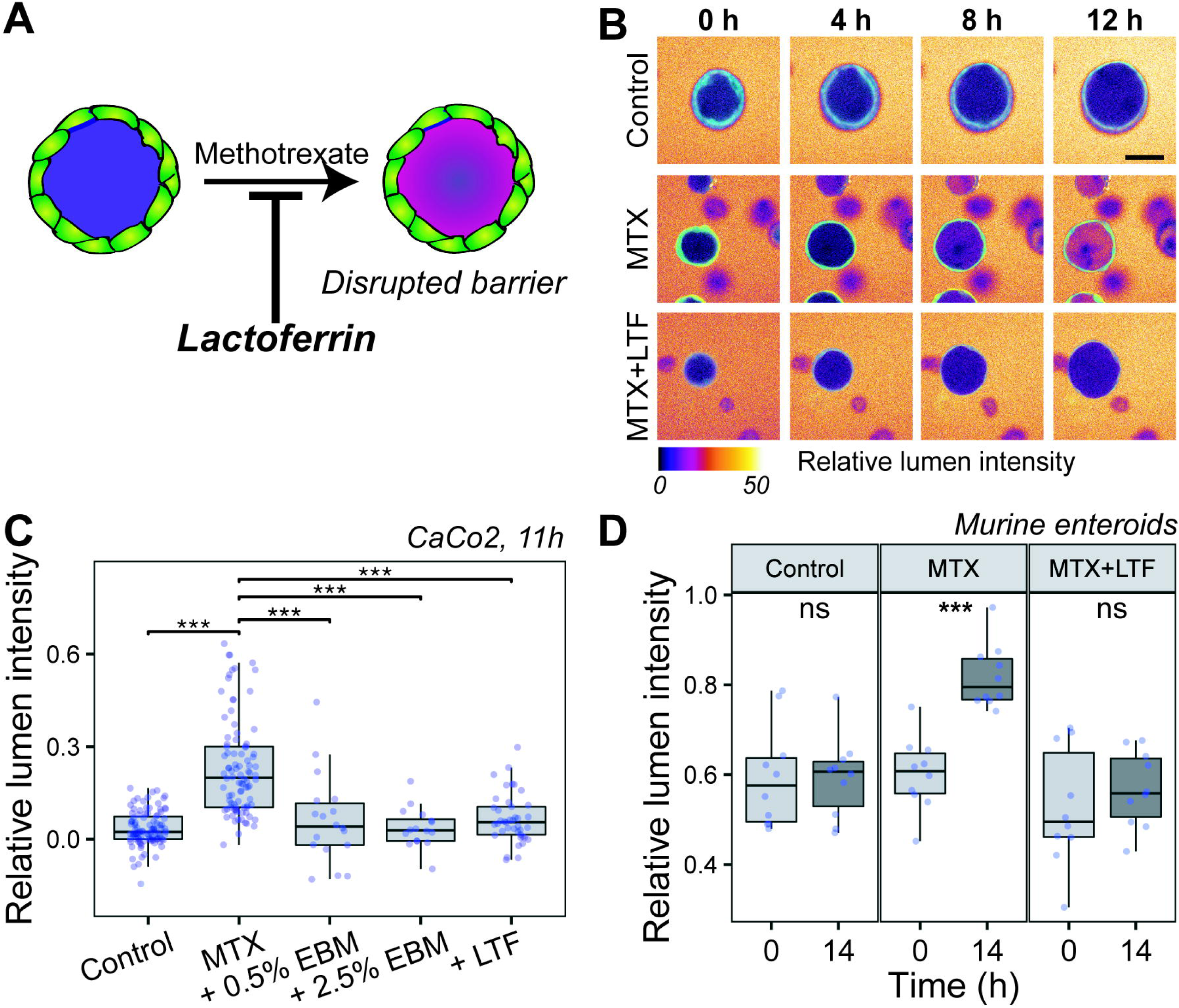
Lactoferrin blocks methotrexate-induced barrier dysfunction in organoids and Caco2 cysts. **A.** A schematic of the experimental model illustrating increased luminal fluorescence upon induction of barrier dysfunction. **B.** Fluorescence micrograph time series of Caco2 spheroids upon exposure to 220 nM methotrexate alone (**MTX**) or with 10 μM lactoferrin (**MTX+LTF**). Scale bar: 50µm **C.** Boxplots representing the relative change in intraluminal fluorescence intensity in Caco2 spheroids after 11 h of treatment, due to flux of a 3 kD texas-red labeled dextran into lumen after exposure to methotrexate (220 nM), methotrexate and lactoferrin (10 µM), or methotrexate and expressed breast milk (EBM) (either 0.5% or 2.5% of growth media). Each dot represents an individual spheroid. Statistical significance is based on non-parametric Wilcox test compared to methotrexate treatment. **D.** Boxplots representing the change in relative fluorescence due to flux of 3 kDa cascade blue-labeled dextran into the lumen of Murine Enteroids upon methotrexate or methotrexate-lactoferrin exposure. Each dot represents an individual enteroid. Statistical significance is based on non-parametric Wilcox test compared to time = 0 h.

We next investigated the specific molecular activity responsible for the effects of expressed breast milk (EBM) on epithelial barrier function. Lactoferrin is present at a concentration of 13-113 µM in breast milk^17^, and it has been hypothesized that it is one of the factors responsible for its barrier-protective effects^41^. We therefore tested whether lactoferrin was sufficient to restore barrier integrity in this assay by titrating cysts with increasing concentrations of human lactoferrin (Supplemental Figure S1C) in combination with methotrexate. We noted a dose-dependent protective effect that saturated at higher lactoferrin concentrations (EC50 = 6.5 µM), with no significant effect noted for concentrations below 1 µM. Therefore, for subsequent experiments, we used 10 µM lactoferrin, a concentration that consistently blocked luminal accumulation of 3 kDa dextran in methotrexate-treated cysts (Figure 1B, C, Supplemental Figure S1A, B). We also evaluated the most common human milk oligosaccharides (HMOs), 2-Fucosyllactose (2-FL) and 3-Fucosyllactose (3-FL), in this assay and found that these components had no significant effect on epithelial barrier function, consistent with previous studies suggesting they function primarily as prebiotics (Supplemental Figure S2A)^42^. Similar results were obtained in the higher complexity murine intestinal epithelial organoid model^30^. Treatment of organoids with methotrexate after 3 days in culture resulted in loss of barrier function, which was restored upon treatment with 10 µM lactoferrin (Figure 1D, Supplemental figure S1D). We also assessed whether lactoferrin could reverse barrier dysfunction (as opposed to blocking initiation). We again exposed Caco2 cysts to methotrexate, but first for four hours to induce barrier dysfunction, after which methotrexate was removed and lactoferrin was added. Again, we observed accelerated barrier function improvement following lactoferrin administration (Supplemental Figure S1E).

### Methotrexate Induces Activation of an Epithelial-Mesenchymal Transition (EMT) Program

To identify the molecular drivers for methotrexate-induced barrier dysfunction, we conducted RNA sequencing on Caco2 spheroids treated with methotrexate and lactoferrin. Given our observation that barrier function was compromised as early as 4-6 hours after exposure to methotrexate, we isolated RNA from spheroids 4-hour post-treatment to capture the earliest transcriptional events responsible for initiating the processes. Differential gene expression analysis revealed major transcriptional changes upon methotrexate exposure, which were mostly normalized upon lactoferrin co-treatment (Figure 2A). Gene ontology analysis noted the activation of an EMT program^43^ in methotrexate treated cells that was absent in controls or lactoferrin treated cells (Figure 2B). Transcriptional changes included upregulation of SNAI2, TWIST1, ZEB2, ZEB1, N-cadherin, and STAT3, as well as downregulation of E-cadherin and Occludin (Figure 2C, **Supplemental Table 2**). Changes in expression of these genes aligns with both the genes upregulated in initiating EMT as well as the alterations to tight and adherens junctions during EMT. We also observed upregulation of lactoferrin transcripts in MTX exposed cells, consistent with lactoferrin’s role as a signaling molecule involved in inflammation. Phenotypic changes consistent with EMT were also noted in organoid culture by immunofluorescence, including a reversal of epithelial polarity and redistribution of actin (Figure 3A, B).

**Figure 2:**
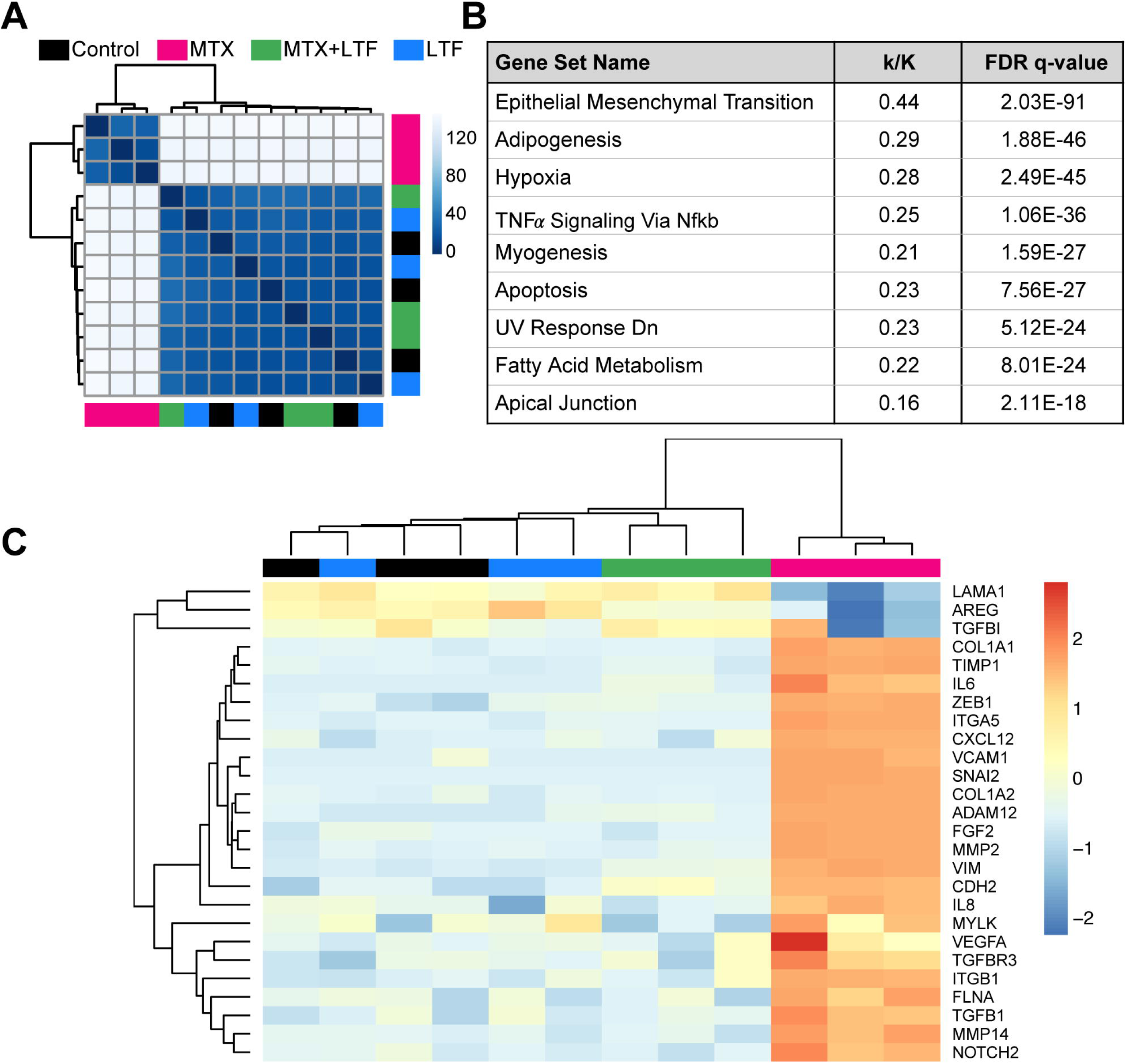
Methotrexate induces and EMT transcriptional program in Caco2 cysts that is reversed by lactoferrin. **A.** matrix representing Euclidean distance between samples based on the rlog-normalized counts from bulk RNAseq data demonstrating transcriptional differences across the various exposure conditions. **B.** Gene sets enriched in the top 500 genes upregulated upon methotrexate treatment in Caco2 spheroids. **C.** Heatmap showing changes in gene expression for a subset of genes. High level of activation of EMT specific genes such as ZEB1, SNAI2, and VCAM1, as well as TGFB1 is seen after exposure to methotrexate and normalized upon addition of lactoferrin.

**Figure 3:**
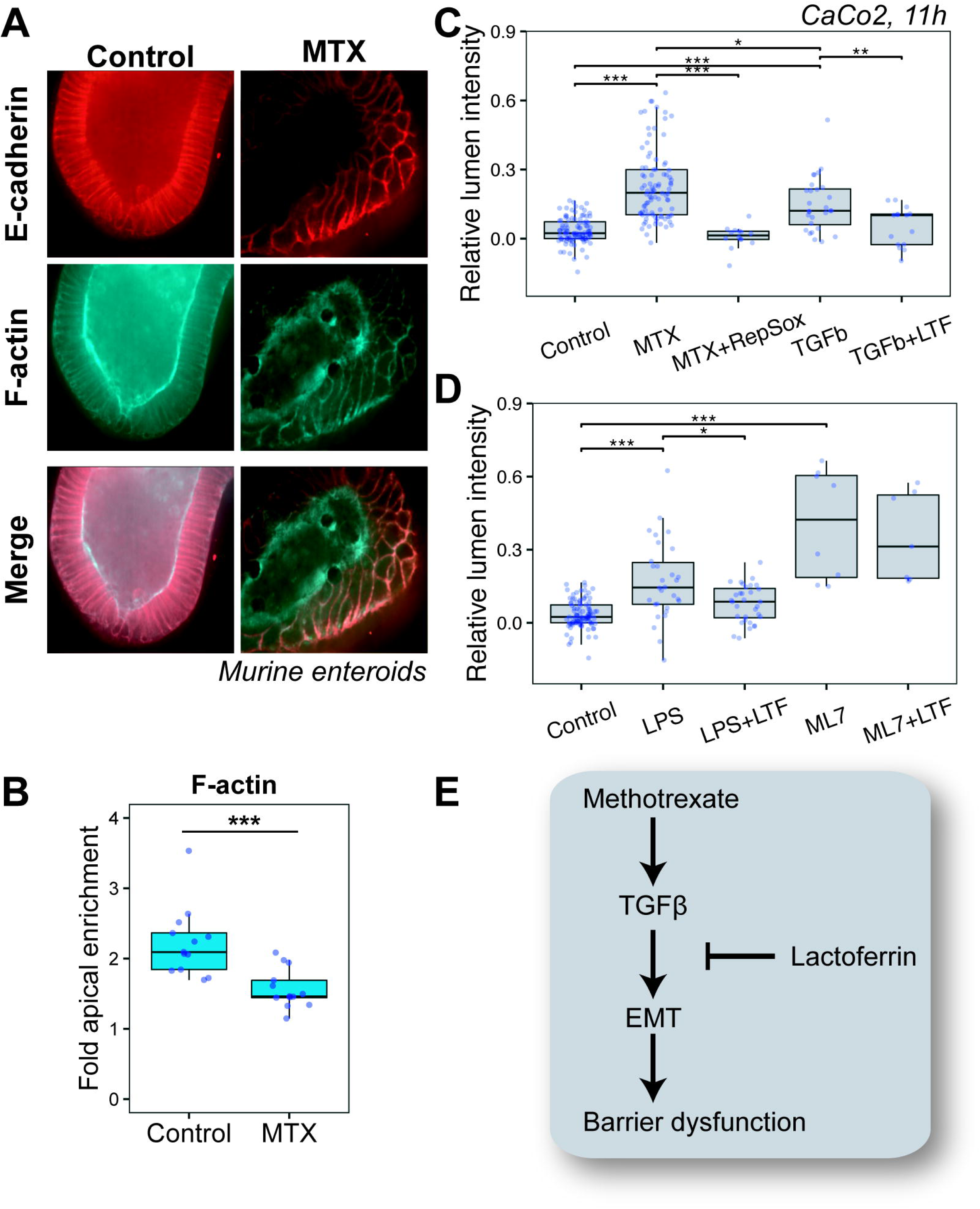
Lactoferrin blocks TGF-β signaling to reverse EMT in Caco2 cysts. **A.** E-cadherin and F-actin immunofluorescence staining in murine organoids after exposure to methotrexate reveal redistribution of Actin and E-cadherin in murine organoids after exposure to methotrexate. **B.** Fold-change in the apical phalloidin intensity of the organoids after exposure to methotrexate. **C.** Relative luminal fluorescence intensity of Caco2 spheroids after incubation with methotrexate or TGF-β (to induce barrier dysfunction), and RepSox or lactoferrin (to inhibit TGF-β signaling). **D.** Box-plot representing the relative luminal fluorescence in LPS, LPS-LTF, ML7, and ML7-LTF exposure conditions. Each dot represents the individual spheroids. Statistical significance is based on non-parametric Wilcox test compared to control or methotrexate treatment. **E.** Schematic illustrating the proposed mechanism of lactoferrin blockade of EMT in Caco2 cysts and murine enteroids.

### Lactoferrin Reverses Barrier Dysfunction by Blocking EMT Downstream of TGF-**β**

Among the observed transcriptional changes induced by methotrexate, we observed a significant upregulation of TGFB1 transcripts (2-fold, p = 0 4.88E-15), (**Supplementary Table 2**), suggesting a potential mechanism for methotrexate-induced barrier dysfunction. Specifically, we reasoned that methotrexate-induced EMT and barrier dysfunction requires activation of a TGF-β autocrine loop, and that lactoferrin could block EMT by inhibiting TGF-β signaling. Consistent with this putative mechanism, prior studies demonstrated that lactoferrin can bind to TGFBR1^15, 44^ suggesting an ability to modulate signaling through this pathways.

To first test the hypothesis that signaling through the TGF-β pathway is necessary for methotrexate-induced barrier dysfunction, we repeated the barrier permeability assay in the presence of RepSox, an ATP-competitive inhibitor of the ALK5 subunit of TGFBR1 receptor^45^. RepSox administered at 100 ng/ml (348 nM) restored barrier function similar to lactoferrin (Figure 3C, Supplemental Figure S2B). To next test the hypothesis that aberrant TGF-β signaling is sufficient to induce barrier dysfunction in this assay, we treated Caco2 cysts with 16 nM (200 ng/ml) TGF-β and found that it phenocopied the effects of methotrexate treatment on the epithelium. Finally, to determine whether Lactoferrin was acting to specifically disrupt TGF-β signaling, we repeated the assay in the presence of 16 nM TGF-β and 10 μM lactoferrin (Figure 3C). We again observed a complete block of TGF-β induced barrier dysfunction. Together, these data demonstrate that TGF-β signaling is necessary and sufficient to induce barrier dysfunction downstream of methotrexate, and are consistent with a model wherein lactoferrin blocks EMT-mediated barrier dysfunction by inhibiting autocrine TGF-β signaling.

To investigate the generalizability of lactoferrin’s barrier protective effect, we repeated the assay using several other agents known to induce barrier dysfunction. LPS is a byproduct of gram negative bacteria common to the intestine and similar to methotrexate, induces EMT^46^ as well as barrier dysfunction. Exposure of LPS at a concentration of 0.016 nM (40 ng/ml) led to decreased barrier function, which was restored by 48% by concurrent exposure to 10 μM lactoferrin (Figure 3D). Exposure of Caco2 spheroids to antagonists of cellular contractility, such as the myosin light chain kinase inhibitor ML7, also induced rapid barrier dysfunction at 10 µM^47^. However, ML-7 induced barrier dysfunction was not reversed by exposure to lactoferrin (Figure 3D, Supplemental Figure S2C). Together, these data suggest that lactoferrin selectively restores barrier function induced by pathways downstream of the TGF-β (Figure 3E).

## DISCUSSION

To identify the mechanism of lactoferrin’s impact on intestinal permeability induced by methotrexate and other compounds, we developed an *in vitro* model devoid of a complex microbiota and immune cells. This assay allowed us to specifically interrogate the activity of lactoferrin and other components of breast milk on the epithelial compartment. Our data suggest that human holo-lactoferrin can act to restore barrier function downstream of methotrexate treatment by inhibiting activation of an EMT program in intestinal epithelial cells (murine organoids and Caco-2 cells). We further identify a mechanism for EMT blockade by lactoferrin-mediated inhibition of TGF-β signaling. Lactoferrin restored barrier function even after initial sustained disruption with methotrexate, a finding consistent with interruption of a TGF-β-mediated feedback loop which sustains EMT^48^. We additionally demonstrate that this barrier protective effect is generalizable to other EMT-inducing compounds, as lactoferrin protects against agents such as LPS and TGF-β. In contrast, lactoferrin does not protect against mechanistically distinct disrupters of barrier function such as ML7, which acts by inhibiting epithelial contractility.

Lactoferrin is secreted by a wide variety of cells types as an inflammatory mediator *in vivo*. It serves as a transcriptional regulator, entering the cell through its dedicated receptor (LfR^16^). Lactoferrin has also been shown to bind to TGF-β receptors and to stimulate TGF-β pathways in both immune cells and osteoblasts^15, 44^. In contrast, we find that lactoferrin also has activity as an inhibitor of TGF-β signaling and methotrexate induced EMT. Given the capacity of lactoferrin to interact with TGF-β receptors, it may be acting as a competitive inhibitor of TGF-β binding (e.g. a partial agonist). Further investigation will be necessary to interrogate these more detailed aspects of lactoferrin’s activity on the TGF-β pathway.

Methotrexate is one of the most widely used anti-inflammatory and chemotherapeutic agents in the world. The primary reason for discontinuation or complication of therapy are secondary GI effects, including pain, severe mucositis, and hepatic injury^7, 9^. As it is known that methotrexate causes intestinal epithelial barrier dysfunction even at low dosages^8^, it is likely that much of the secondary effects are driven by this barrier disruption. Coadministration of oral lactoferrin may allow for a protective effect in the gut without impeding anti-inflammatory effect in patients with diseases such as rheumatoid arthritis, or chemotherapeutic effects. While further study is necessary, it is likely that a gut-protective effect could be achieved without impact to therapeutic effect on other anatomical locations or circulating cells, preserving anti-inflammatory and chemotherapeutic function while simultaneously improving patient tolerance and secondary complications.

Our *in vitro* findings are consistent with previous *in vitro* and *in vivo* findings. Bovine lactoferrin has previously been noted to inhibit EMT in the context of squamous cell carcinoma cells^49^, demonstrating a broad ability to inhibit EMT in multiple epithelial cell types. This is further reinforced by studies examining the effect of lactoferrin on wound healing in both skin^50^ and corneal tissue^51^ in model organisms. These studies demonstrate the ability of lactoferrin to promote keratinocyte proliferation while simultaneously driving re-epithelialization, a finding which is phenotypically congruent with lactoferrin’s ability to inhibit drivers of EMT as shown in our data. From a functional standpoint, we speculate that the value of this effect is to allow for a proliferative response while simultaneously reinforcing epithelial barrier function and decreasing TGF-β mediated fibrosis^52^. This effect is not limited to skin, as lactoferrin has demonstrated the ability to improve intestinal regeneration after injury in rats^53^

Our data also suggest a possible mechanism for observations in clinical studies of lactoferrin supplementation. Trials have demonstrated the capacity of lactoferrin to ameliorate or provide prophylaxis for neonatal enterocolitis^22^ and post-antibiotic diarrhea^25^. NEC is a severe intestinal illness effecting primarily premature infants^54^. The pathophysiology is thought to be intrinsically tied to disruption of barrier function and studies have demonstrated significant upregulation of TGF-β in affected tissues, suggesting a possible role for EMT^55^. Post-antibiotic diarrhea is thought to be mediated by shifts in commensal bacteria which alters barrier function but antibiotics themselves have been shown to have a direct impact on barrier function^56^. Not only does our data demonstrate possible therapeutic potential in disease, it also suggests a strong potential for application of lactoferrin as an agent capable of ameliorating methotrexate induced side effects in the clinic. By inhibiting methotrexate-driven TGF-β activation and preventing EMT, lactoferrin may protect from side effects such as abdominal pain, mucositis, and increased risk of infection secondary to bacterial translocation. By inhibiting this process lactoferrin may allow for both improved safety in chemotherapeutic protocols which utilize methotrexate, as well as improving tolerance of methotrexate given for therapy of autoimmune disease such as inflammatory bowel disease or rheumatoid arthritis. Encouragingly, recent evidence suggest that lactoferrin can be formulated for efficient delivery to the small intestine^57^. While further work remains necessary to establish generalizability of this effect, it also remains possible that lactoferrin may be of assistance in mitigating side effects of other medications.

---

**Supplementary Figure 1:**
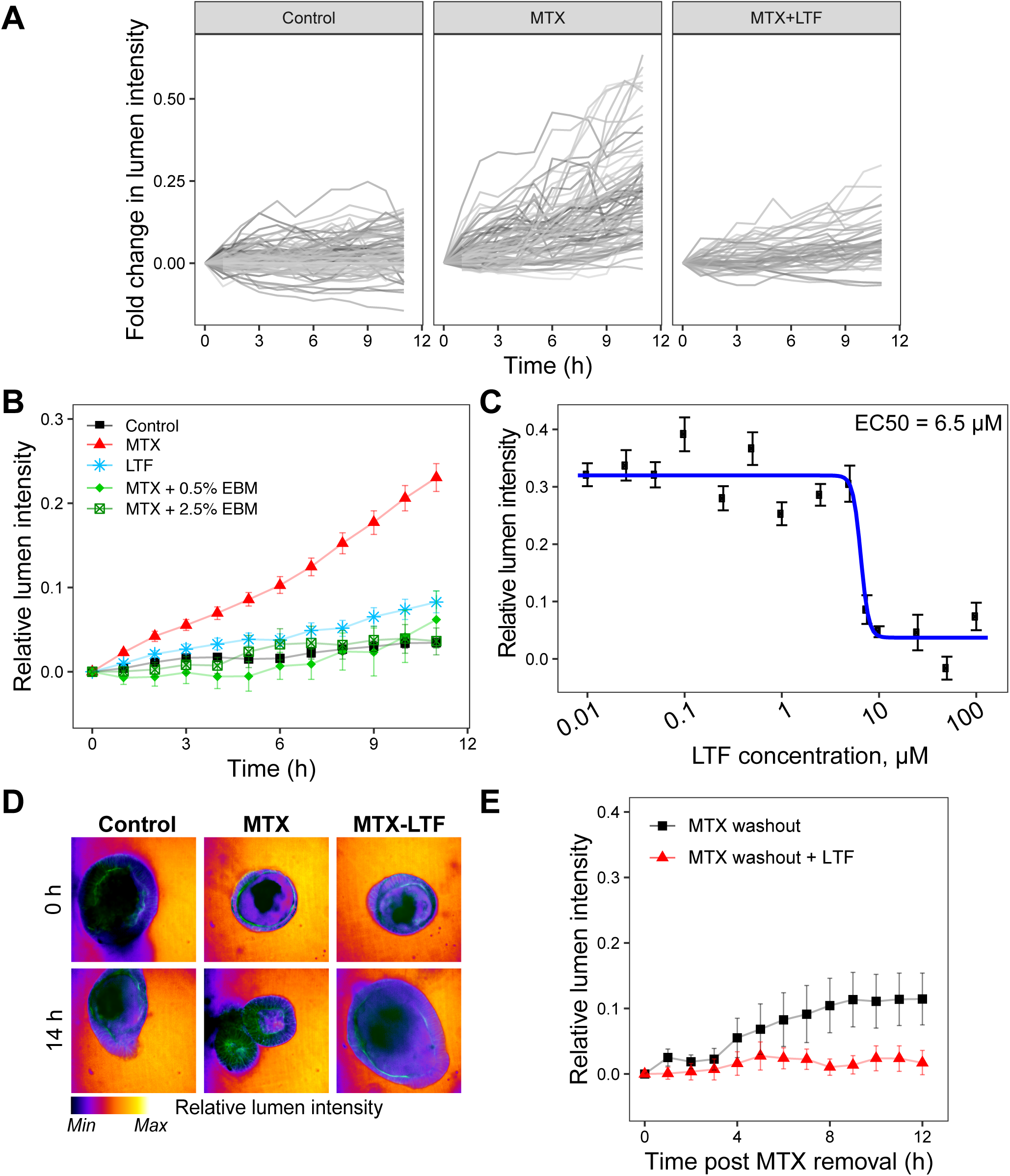
**A.** Distribution of relative luminal fluorescence over time across individual spheroids. **B.** Relative luminal fluorescence over time for Caco2 spheroids upon exposure to methotrexate (220 nM), lactoferrin (10 µM), methotrexate and lactoferrin, or methotrexate and either 0.5% expressed breast milk or 2.5% expressed breast milk (EBM) mixed with standard growth media. **C.** Dose finding curve for lactoferrin’s protective effect from methotrexate (220 nM)-induced barrier disruption. **D.** Pre- and post-exposure images of murine organoids exposed to methotrexate (220 nM) or methotrexate and lactoferrin (10 µM) demonstrating barrier disruption of methotrexate and protective effect of lactoferrin. **E.** Relative luminal fluorescence as a function of time in Caco2 spheroids upon methotrexate-washout after an initial 4 h methotrexate exposure, with or without lactoferrin exposure.

**Supplementary Figure 2:**
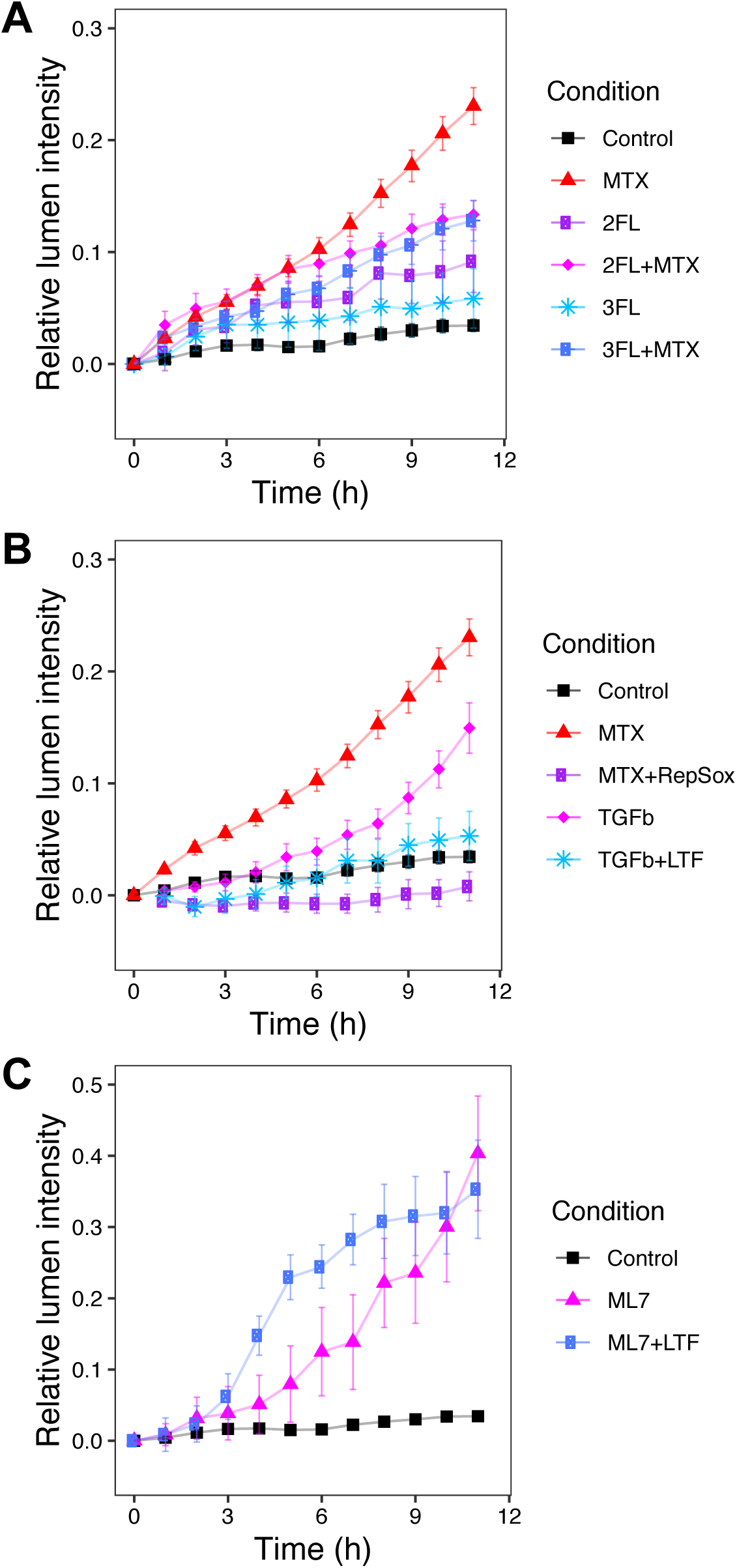
**A.** Relative luminal fluorescence over time for Caco2 spheroids after exposure to methotrexate (220 nM), Methotrexate + RepSox (348 nM), TGF-β (16 nM), or TGF-β and Lactoferrin (10 µM) **B.** Relative luminal fluorescence over time for Caco2 spheroids after exposure to ML7 (10 µM) or ML7 and Lactoferrin (10 µM) **C.** Relative luminal fluorescence over time for Caco2 spheroids after exposure to Methotrexate (220 nM), 2-Fucosyllactose (2-FL) (400 nM), 3-Fucosyllactose (3-FL) (150 nM), or Methotrexate and 2-FL or 3-FL, demonstrating possible protective effect but no statistically significant findings.

